# Revealing the spatial characteristics of rice heat exposure in Japan through panicle temperature analysis

**DOI:** 10.1101/2023.12.26.573387

**Authors:** Yusuke Toda, Yasushi Ishigooka, Mayumi Yoshimoto, Takahiro Takimoto, Tsuneo Kuwagata, David Makowski, Toshihiro Hasegawa

## Abstract

Elevated temperatures during the flowing stage contribute to heat-induced spikelet sterility in rice, posing a major threat to production considering climate change projections. Developing effective strategies for stable rice production through breeding and crop management is critical; however, our understanding of regional, seasonal, and long-term trends in rice heat exposure remains limited. Previous studies on spikelet sterility revealed that panicle temperature, estimated using a micrometeorological model and common meteorological factors, serves as a reliable indicator of rice heat exposure. In this study, we employed this model to identify the differences between panicle and air temperatures (DPAT) and their causes over the past 45 years in Japan. A gridded daily meteorological dataset covering Japan was interpolated at an hourly time step and used as input data of the micrometeorology model for estimating panicle temperatures during flowering. Statistical analysis of the resulting data revealed an increasing trend in the frequency of rice panicle heat exposure over time across many locations in Japan. During heat-receptive periods, panicle temperature generally exceeded air temperature, indicating the inadequacy of relying solely on air temperature to gauge rice heat stress. DPAT values showed substantial inter-regional variations in both mean values (from −0.5℃ to 3.0℃) and seasonality. Through machine learning and statistical methods, the relationship between DPAT and meteorological factors was characterized, delineating the effects of the meteorological factors on regional and seasonal DPAT variations. Focusing on major high-risk regions, we show that mitigation strategies should be adapted to consider regional characteristics and avoid high DPAT conditions during rice heading periods.

## Introduction

Rice, a staple crop in Japan, is cultivated nationwide under diverse climatic conditions. The approximate growing region spans from 24 °N to 45 °N and from 0 m to >1000 m above sea level. Although most rice-growing regions experience temperate climates, the growing seasons may still be threatened by heat during critical growth stages. In recent years, summers with elevated temperatures have induced spikelet sterility in field conditions (Yoshimoto et al., 2018). This sterility, resulting from poor anther dehiscence due to heat stress during flowering (Satake and Yoshida, 1978; Matsui et al., 2000), directly affects rice yield. With climate change increasing the frequency of heat stress on rice, spikelet sterility is expected to become more prevalent in the future (Shah et al., 2011; Devkota et al., 2013).

Substantial efforts have been made to mitigate heat-induced spikelet sterility through both breeding and crop management. To aid breeders, researchers have sought loci associated with heat tolerance (Ishimaru et al., 2016), facilitating the development of tolerant cultivars through marker-assisted selection. For farmers, adjusting transplanting dates has proven effective in reducing heat-induced spikelet sterility and stabilizing rice production (Ishigooka et al., 2017; Ishigooka et al., 2021).

As a foundational aspect of these studies, identifying regions at high risk of spikelet sterility is crucial for developing and implementing region-specific countermeasures. Risk generally comprises three factors: hazard, exposure, and vulnerability. Here, hazard refers to the increasing heat effect on rice spikelets throughout Japan. Quantifying heat exposure is essential for determining the requisite heat tolerance of future cultivars and establishing future crop management practices. Consequently, understanding the spatial and temporal distribution of rice heat exposure is vital for mitigating future risks.

The regional characteristics of rice heat exposure during critical growth stages remain unclear in Japan, owing to the complex relationship between heat-induced spikelet sterility and meteorological factors. Previous studies have revealed that sterility is induced not only by high air temperature but also by high humidity, low wind speed, and intense solar radiation (Matsui et al., 1997; Matsui et al., 2007; Tian et al., 2010; Hasegawa et al., 2011). The important implication is that regions currently experiencing high temperatures may not necessarily be prone to heat stress leading to sterility, and vice versa for cooler regions. To identify regions at high risk of heat-induced spikelet sterility, a deeper understanding of factors contributing to frequent temperature spikes in panicles is necessary.

Previous studies have indicated that rice panicle temperature more directly reflects spikelet heat stress and serves as a better indicator of spikelet sterility (Yoshimoto et al., 2011; Julia and Dingkuhn, 2013; Oort et al. 2014). Panicle temperature acts as a mediating factor between meteorological conditions and spikelet sterility, suggesting its utility for assessing heat exposure. Given that ample meteorological knowledge about air temperature already exists, accurately estimating the difference between panicle and air temperature (DPAT) is essential to determine regional heat exposure. It has been reported that DPAT can vary by as much as 6°C–7°C depending on environmental conditions (Matsui et al., 2007). Revealing how, when, and where DPAT spikes will enable the evaluation and prediction of sterility risk, complementing existing knowledge regarding air temperature.

To estimate panicle temperature and DPAT, the Integrated Micrometeorology Model for Panicle and Canopy Temperature (IM^2^PACT; Yoshimoto et al., 2011) can be employed. This model, accounting for heat balance around panicles using meteorological variables, has been applied to field survey data in Japan (Yoshimoto et al., 2021) and worldwide (Yoshimoto et al., 2022), proving useful for explaining the occurrence of heat-induced spikelet sterility (Yoshimoto et al., 2021). IM^2^PACT requires only general meteorological factors as inputs, allowing the estimation of panicle temperatures’ spatial distribution using high-resolution meteorological data, such as daily meteorological data obtained from a 1 km grid in Japan for the past 45 years, developed by Seino (1993).

In the present study, we aimed to determine regional and seasonal DPAT characteristics and the weather factors influencing them. Despite the wealth of information on air temperature, a quantitative knowledge of DPAT is lacking as to where and to what extent it appears and how its regional characteristics have changed over the past 45 years. By comprehending these spatial and temporal trends in DPAT, we can provide more precise information for crop management, reducing uncertainty in future forecasts. As a prerequisite, we compared the efficacy of evaluating heat exposure using panicle and air temperatures. Specifically, we simulated daily panicle and air temperatures during flowering in Japan over the past 45 years at a 1 km grid resolution using IM^2^PACT. We then quantified DPAT and analyzed their variations spatially and temporally in Japan using machine learning and statistical methods.

## Materials and methods

### Meteorological data and panicle temperature estimation

We used a gridded daily meteorological dataset with third-order resolution (approximately 1 × 1 km) developed by the National Institute for Agro-Environmental Sciences, Japan. This dataset interpolated observed data from Automated Meteorological Data Acquisition System (AMeDAS) stations (Seino, 1993). Daily mean, maximum, and minimum air temperatures (°C, 1.5 m above the ground) and solar radiation (MJ m^−2^) were used from this dataset. Additionally, relative humidity (%), precipitation (mm), and wind speed (m s^−1^, 2.5 m above the ground) were prepared in the same manner (Ishigooka et al., 2017). Furthermore, “Land Use Mesh 2006” (http://www.mlit.go.jp/kokudokeikaku/gis/index.html), provided by the Ministry of Land, Infrastructure, Transport and Tourism of Japan, was linked with each grid cell (Ishigooka et al., 2011). Grid cells containing paddy fields covering >1% of the grid area were selected for analysis. The resulting meteorological data comprised 7 climate variables, 152,697 grid cells, 45 years (1978–2022), and 153 days (June 1–October 31).

Hourly panicle temperature was estimated using these data. After estimating hourly meteorological factor values using trigonometric functions (Maruyama et al., 2017), IM^2^PACT (Yoshimoto et al., 2011) was employed to generate hourly panicle temperature. The resulting panicle temperature from 09:00 to 15:00, a typical flowering time for Koshihikari, a common Japanese rice cultivar (Ishimaru et al., 2010), was averaged. Air temperature during the flowering period was calculated using the same method for comparison, based on hourly air temperature estimates obtained via artificial interpolation. The resulting air and panicle temperatures during flowering time were used to compute the temperature differences (DPAT = panicle temperature − air temperature) for each year and grid cell.

### Air and panicle temperatures, and heat exposure around the heading date

Initially, we compared the spatial distribution of air and panicle temperatures around the heading date to identify the more suitable indicator of rice heat exposure. Heading dates were obtained for each subadministrative region from a dataset provided by the Ministry of Agriculture, Forestry, and Fisheries, Japan, following the approach of Ishigooka et al. (2017). These dates were allocated to each grid cell (Ishigooka et al., 2011).

Next, we analyzed the long-term trend in rice heat exposure using panicle temperature. Heat exposure frequency (HEF) was defined as the ratio of days with heat exposure around the heading date for each year and grid cell. We targeted 15 days around heading, accounting for variations in heading dates within ±1 week due to individual differences or crop management. To determine the occurrence of heat exposure, we applied the threshold of 33°C for panicle temperature, as it was reported by Yoshimoto et al. (2021) as the average panicle temperature during flowering time, with rice spikelet sterility increasing significantly at higher temperatures. The time trend for HEF was statistically tested for each grid cell using a quasibinomial generalized linear model through the R function “glm.”

Finally, we identified several regions with high frequencies of heat stress. These regions were defined by identifying subadministrative regions and selecting grid cells where the mean HEF in 2008–2022 exceeded 15%.

### Analysis of mean DPAT

To focus on spatial variation, DPAT and meteorological data were averaged over days and years for each grid cell. Six meteorological factors, as outlined in the “Meteorological data and panicle temperature estimation” section, were used, excluding daily mean temperature. DPAT values during flowering time were averaged, whereas daily values were averaged for meteorological factors. A random forest machine learning model (Breiman, 2001) was trained on these data to quantify the effects of meteorological factors on DPAT spatial variability. Partial dependence plots were generated using the trained model to illustrate the mean DPAT response to each meteorological variable. Shapley values were also computed to identify the most influential meteorological variables. Shapley values, rooted in game theory, are quantitative evaluations of each explanatory variable’s contribution to target variable prediction. An individual Shapley value was calculated for each model input and each individual prediction. For a given model prediction, the sum of the Shapley values obtained for all model inputs was equal to the difference between the prediction under consideration and the mean value of the predictions derived from the dataset. Positive or negative Shapley values indicated whether the considered explanatory variable contributed to an increase or decrease in predicted DPAT.

The “randomForest” R package (ver. 4.7-1.1; Liaw et al., 2002) was used to model the relationship between the temporal means of meteorological variables and DPAT., Hyperparameters “mtry” and “ntree” were selected from mtry = (2, 3, 4, and 5) and ntree = (100, 200, 300, 400, and 500) using threefold cross-validation. In this case, mtry = 4 and ntree = 500 were found to minimize root mean squared errors (Table S1). The R package “iml” (ver. 0.11.1; Molnar et al., 2018) was employed to create partial dependence plots and compute Shapley values.

### Analysis of seasonal change in DPAT

To evaluate variation in DPAT within a year and regional differences, seasonal changes in DPAT and meteorological factors were analyzed. All time-series for each variable in each grid cell, spanning 153 days and 45 years, were decomposed into long-term trends, seasonal changes, and residuals using the R function “decompose.” Daily values (153) representing seasonal changes for each grid cell were obtained. Principal component analysis (PCA) was then applied to the estimated seasonal changes of all grid cells to visualize regional variation in DPAT seasonal change. Furthermore, we explored which meteorological factor predominately influenced regional variation. Regression analysis was used to evaluate the effects of meteorological factors on the first and second principal components (PCs). Monthly means of seasonal changes for each meteorological factor and grid cell served as explanatory variables, and 153 daily values were summarized into 5 variables. Regression coefficients were estimated using ridge regression through the “cv.glmnet” function in the R package “glmnet” (ver. 4.1–6).

## Results

### Distribution of air and panicle temperatures around heading dates

Comparing air and panicle temperatures over 15 years several decades ago (1978–1992) and 15 more recent years (2008–2022), we found that the mean air and panicle temperature around the heading date increased by 1.24℃ (Fig. 1a, b) and 1.13℃ (Fig. 1c, d), respectively. Additionally, a difference in the spatial distributions of air and panicle temperature was revealed, with air temperature exhibiting substantial spatial heterogeneity (Fig. 1b), whereas panicle temperature reached elevated levels over a wide area, except for the northern area (Fig. 1d).

**Figure 1.**
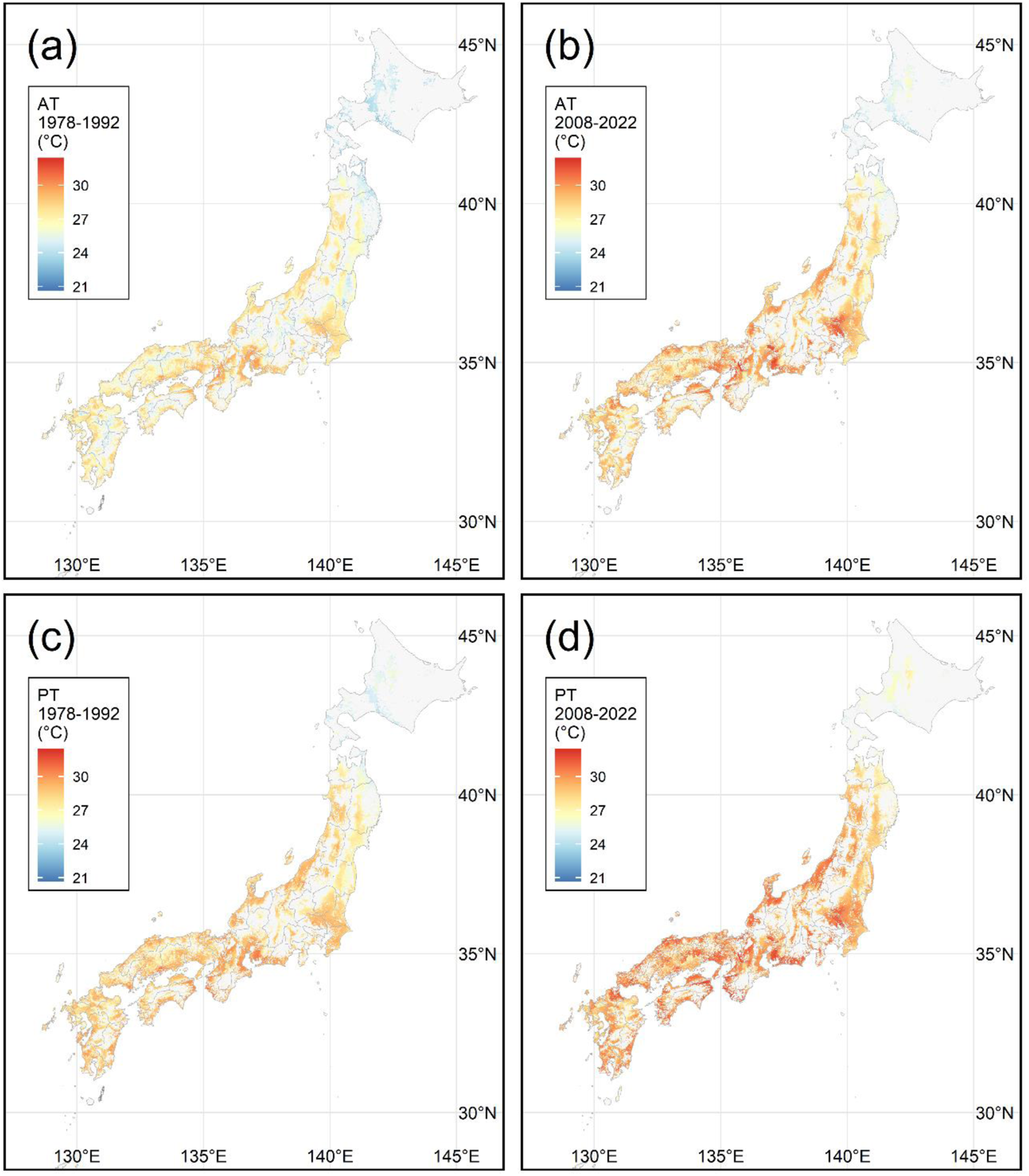
Mean air and panicle temperatures at flowering time during 15 days around heading. Maps show mean air temperatures in (a) 1978–1992 and (b) 2008–2022, and mean panicle temperatures in (c) 1978–1992 and (d) 2008–2022. Mapped values represent mean temperatures over 225 days, i.e., 15 days in 15 years for each grid cell.

### HEF analysis

We compared HEF around heading dates using panicle temperatures and found an increasing trend across a wide area of Japan. The maximum HEF reached 7.07% in 2008– 2022 (Fig. 2a), contrasting with lower HEF in 1978–1992 (Fig. 2b). Statistical tests indicated a significant (*p* < 0.01) yearly effect on HEF in several regions located from the north to the south of Japan (Fig. 2c). Analysis of mean HEF for eight districts weighted according to the rice field area ratio of each grid cell (Fig. 3), incorporating data from 2023 (which included the hottest summer in recorded history), revealed an accelerated rate of increase in recent years, especially in Tohoku, Kanto, and Chubu regions but not in Hokkaido.

**Figure 2.**
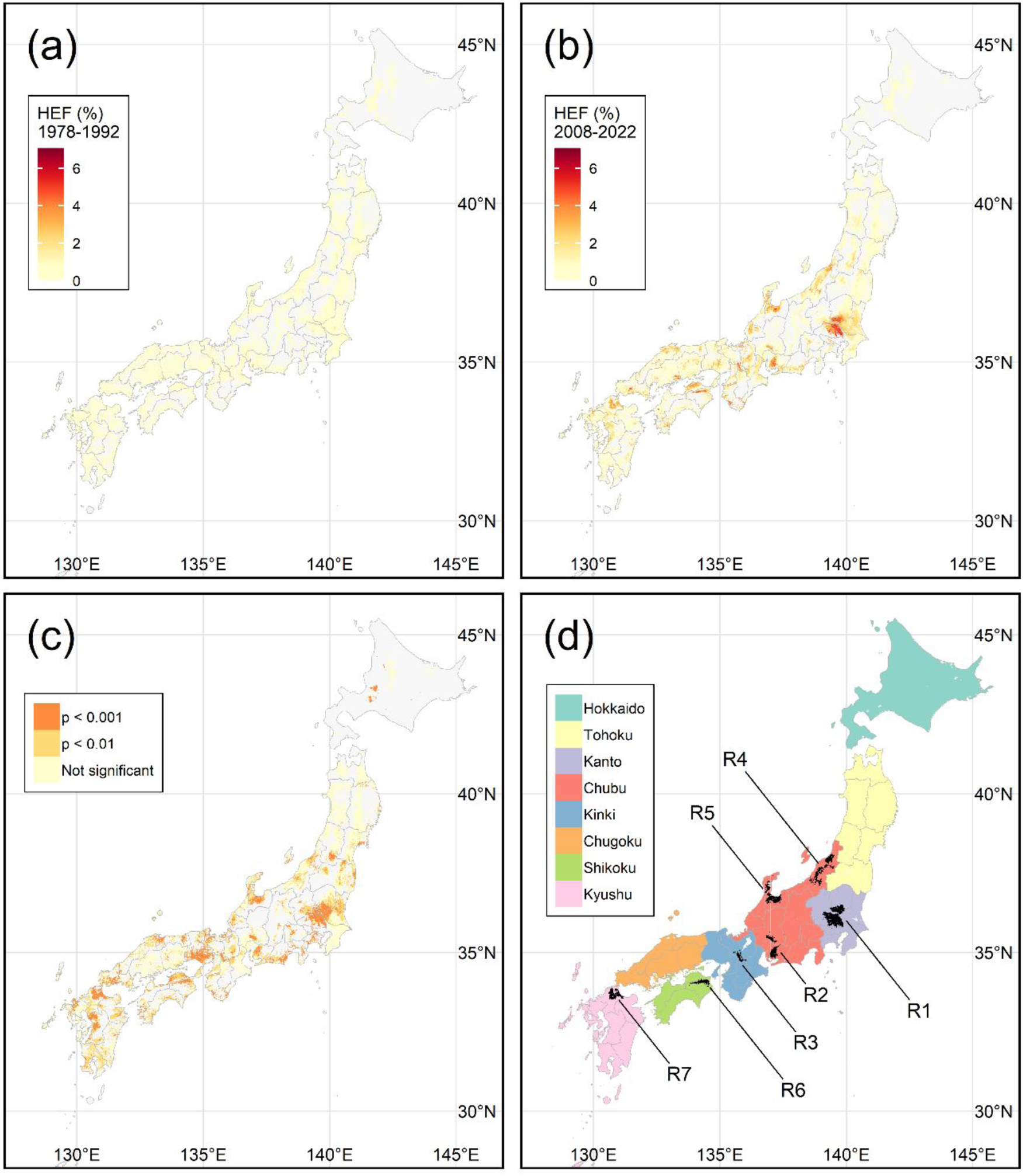
Heat exposure frequency (HEF): the ratio of days with panicle temperature at flowering time exceeding 33℃ during 15 days around heading. (a) HEF in 1978–1992. (b) HEF in 2008–2022. In (a) and (b), HEF is the ratio of the number of days with heat exposure over 225 days (15 days in 15 years). (c) P-values of statistical tests for the increase in HEF in each grid cell. P > 0.01 was considered nonsignificant. (d) Seven regions (R1–R7) identified as high-risk regions with elevated HEF. The eight districts shown in Fig. 3 are color-coded.

**Figure 3.**
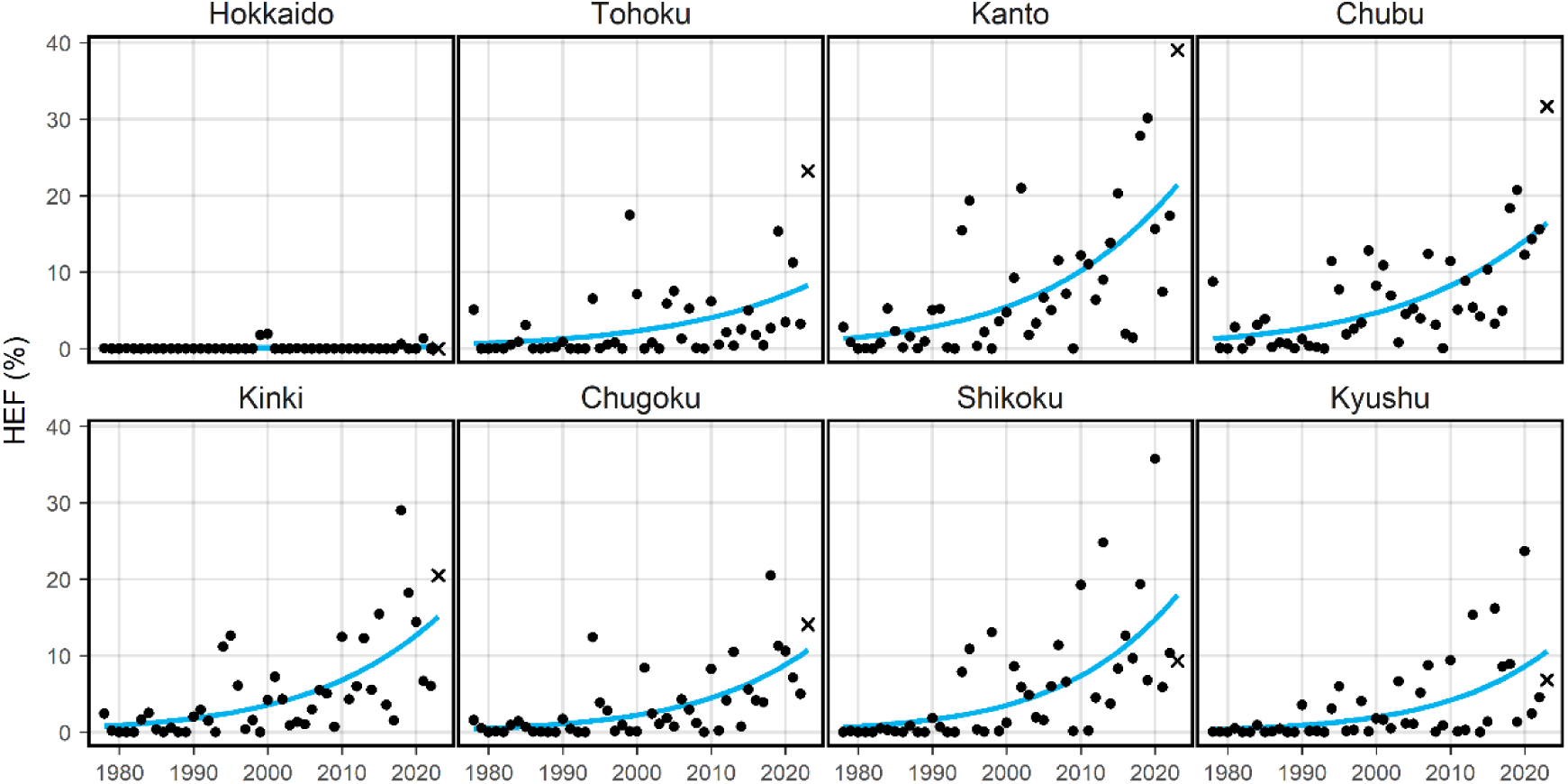
Yearly change in HEF for each district of Japan. Each point corresponds to a mean HEF weighted by the rice field area ratio for each grid cell. Results from 2023 are included and marked with crosses (×). Regression curves obtained using generalized linear models are shown in blue. District locations are shown in Fig. 2d.

We identified seven high-risk regions (R1–R7; Fig. 2d) for focused analysis, including the following subadministrative regions: R1, Saitama Prefecture and Nanbu region in Tochigi Prefecture; R2, Chuno and Touno regions in Gifu Prefecture and Nishimikawa region in Aichi Prefecture; R3, Nanbu region in Kyoto Prefecture; R4, Kaetsukita, Kaetsuminami, Chuetsu, and Uonuma regions in Niigata Prefecture; R5, Toyama Prefecture and Noto region in Ishikawa Prefecture; R6, Hokubu region in Tokushima Prefecture; and R7, Kitakyushu, Chikuzen, and Chikuho regions in Fukuoka Prefecture. Additional information regarding these high-risk regions is provided in Table S2.

### Spatial heterogeneity in mean DPAT

The mean DPAT (panicle temperature − air temperature) exhibited substantial spatial variation, ranging from −0.5℃ to 3.0℃ across different grid cells (Fig. 4b). Overall, DPAT tended to increase with higher elevation (Fig. 4a). The random forest model’s partial dependence plots indicated that low daily maximum temperature, high relative humidity, intense solar radiation, and slow wind speed were the key factors associated with high DPAT values (Fig. 4c). Notably, areas with wind speeds below 2 m s^−1^ correlated high DPAT values (Fig. 4c).

**Figure 4.**
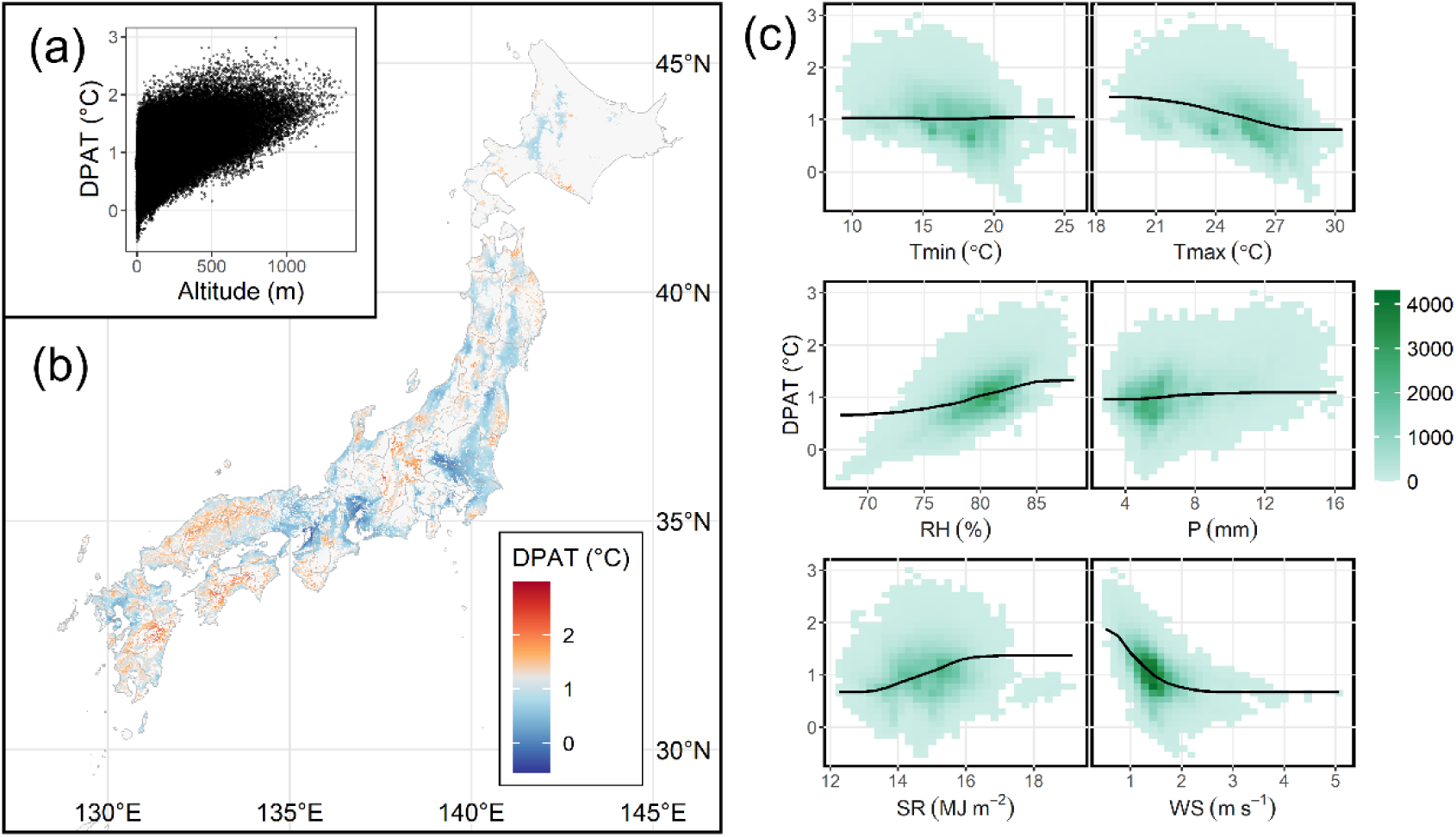
Spatial variation of DPAT, and the relationship between DPAT and meteorological variables. (a) Relationship of altitude and DPAT. (b) Heatmap of the temporal mean of DPAT. (c) Partial dependence plots of DPAT showing the mean response of DPAT to individual meteorological factors: minimum temperature (Tmin), maximum temperature (Tmax), relative humidity (RH), precipitation (P), solar radiation (SR), and wind speed (WS). Green tiles and black lines indicate the distribution of data and the estimated partial dependence plots, respectively.

Focusing on the high-risk regions (Fig. 2d), we found that they could be separated into two groups based on mean DPAT (Fig. 5a): low-DPAT regions (R1–R3) and high-DPAT regions (R4–R7). The effects of meteorological factors on DPAT in these regions were also assessed using Shapley values (Fig. 5b), with differences in these factors characterizing DPAT in each region. In low-DPAT regions, low solar radiation reduced DPAT in R1, whereas high maximum temperature and low relative humidity reduced DPAT in R2 and R3. In high-DPAT regions, high humidity in R4 and R5 and high solar radiation in R6 and R7 were the main factors associated with increased DPAT. Shapley values confirmed that low wind speeds tended to increase DPAT by up to +1°C in all regions (low WS values were associated with positive Shapley values).

**Figure 5.**
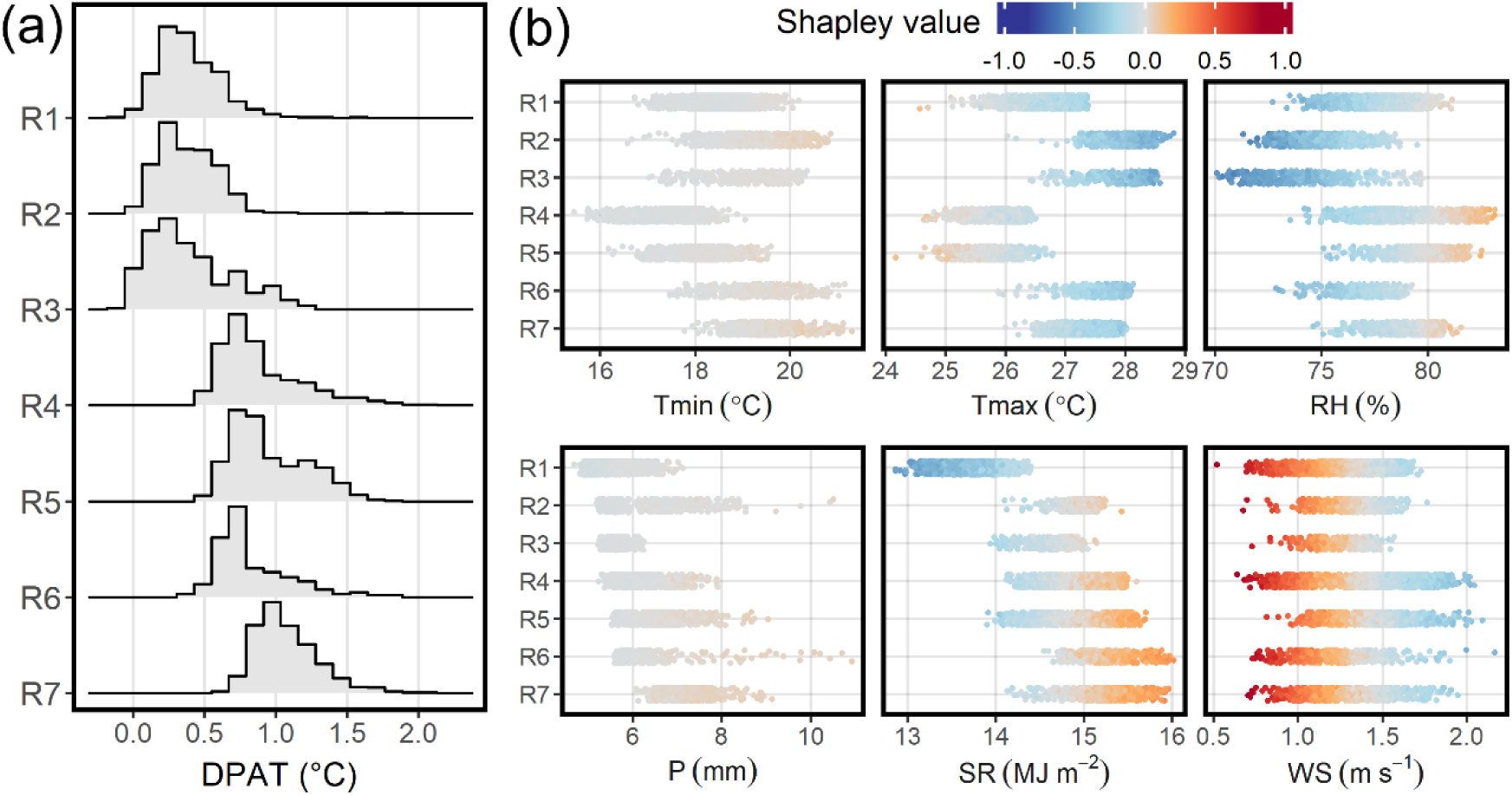
DPAT and Shapley values for each high-risk region. (a) Histogram of DPAT temporal means in each region. (b) Shapley values associated with meteorological variables in each grid cell (indicating the contributions of each factor to predicted DPAT). Colors indicate Shapley values for minimum temperature (Tmin), maximum temperature (Tmax), relative humidity (RH), precipitation (P), solar radiation (SR), wind speed (WS). Positive and negative Shapley values indicate that the corresponding meteorological variables have positive and negative effects on DPAT, respectively.

### Variation in seasonal changes in DPAT

Visualizing the seasonal changes in DPAT (after trend and residues removal) in high-risk regions revealed distinct characteristics (Fig. 6a). Although low-DPAT regions (R1–R3) exhibited small and stable seasonal change, high-DPAT regions (R4–R7) displayed rich seasonal change characteristics, including marked seasonal gradation in R4 and R5, a decrease in September in R6, and a relatively stable trend in R7. These characteristics were also evident in the PCA results, where regions with similar seasonal changes in DPAT showed comparable distributions of the first two PCs (Fig. 6b).

**Figure 6.**
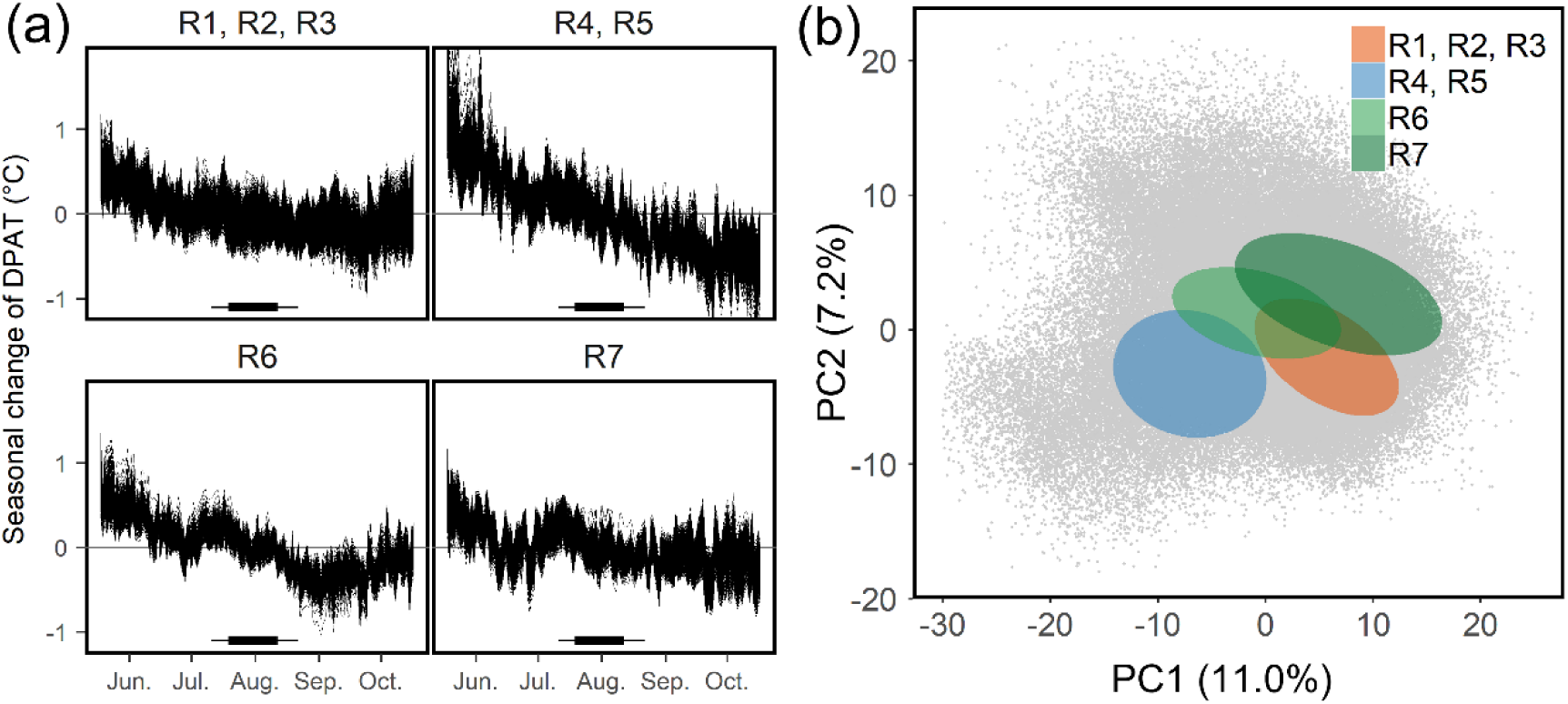
(a) Seasonal changes in DPAT in high-risk regions. Bars at the bottom of each panel indicate the rice heading period, with the thick and thin bars corresponding to 25%– 75% quantile ranges and min–max ranges, respectively. (b) PC score distributions in high-risk regions. Ellipses indicate areas where 90% of points existed, assuming normal distributions for PC scores. Regions with similar PC scores, R1–R3 and R4–R5, were merged. Gray points represent the PC scores of all grid cells. A figure with unmerged regions is also provided (Fig. S1).

We analyzed PC1 and PC2 to determine variations in DPAT seasonal change. The roles of PC1 and PC2 became evident when reproducing seasonal changes with artificial PC scores, focusing on specific PCs while keeping other PCs fixed at zero (Fig. 7a). PC1 represented the linear trend in seasonal change; DPAT rapidly decreased when PC1 was large, whereas DPAT remained stable when PC1 was small. PC2 indicated the location of the peak in seasonal change; the peak occurred in August and June when PC2 was large and small, respectively. Through regression analysis of PCs with the monthly means of meteorological factors (Fig. 7d), we found that solar radiation was the primary driver of both PC1 and PC2. The monthly trend in solar radiation regression coefficients mirrored the variation in seasonal changes represented by PCs. Among other meteorological factors, maximum temperature and wind speed were important determinants of PCs.

**Figure 7.**
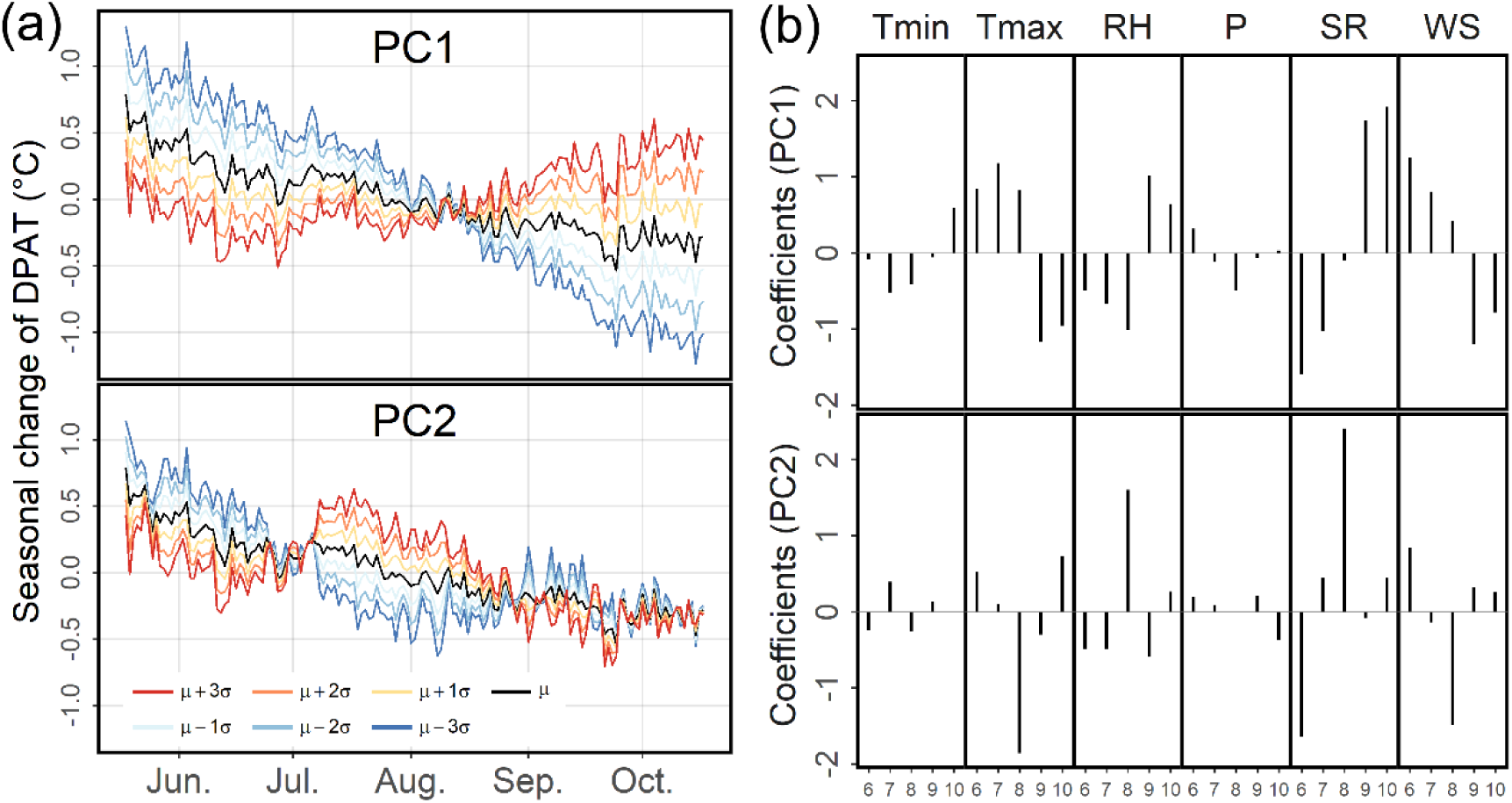
Roles of PCs for seasonal change in DPAT and the meteorological factors affecting this change. (a) Seasonal change in DPAT reproduced using PC1 and PC2. Mean seasonal change is indicated by the black line, and deviations from the mean change when moving PC1 and PC2 ±1, 2, and 3 standard deviations are represented by colored lines. (b) Regression coefficients for PC1 and PC2 on meteorological variables, summarized for each month: Tmin, minimum temperature; Tmax, maximum temperature; RH, relative humidity; P, precipitation; SR, solar radiation; WS, wind speed.

We also visualized the spatial distribution of PC scores (Fig. 8). PC1 scores were related to both latitude and altitude, with PC1 increasing in higher and southern areas and decreasing in lower and northern areas. PC2 was linked to latitude, being higher along the coast facing the Pacific Ocean.

**Figure 8.**
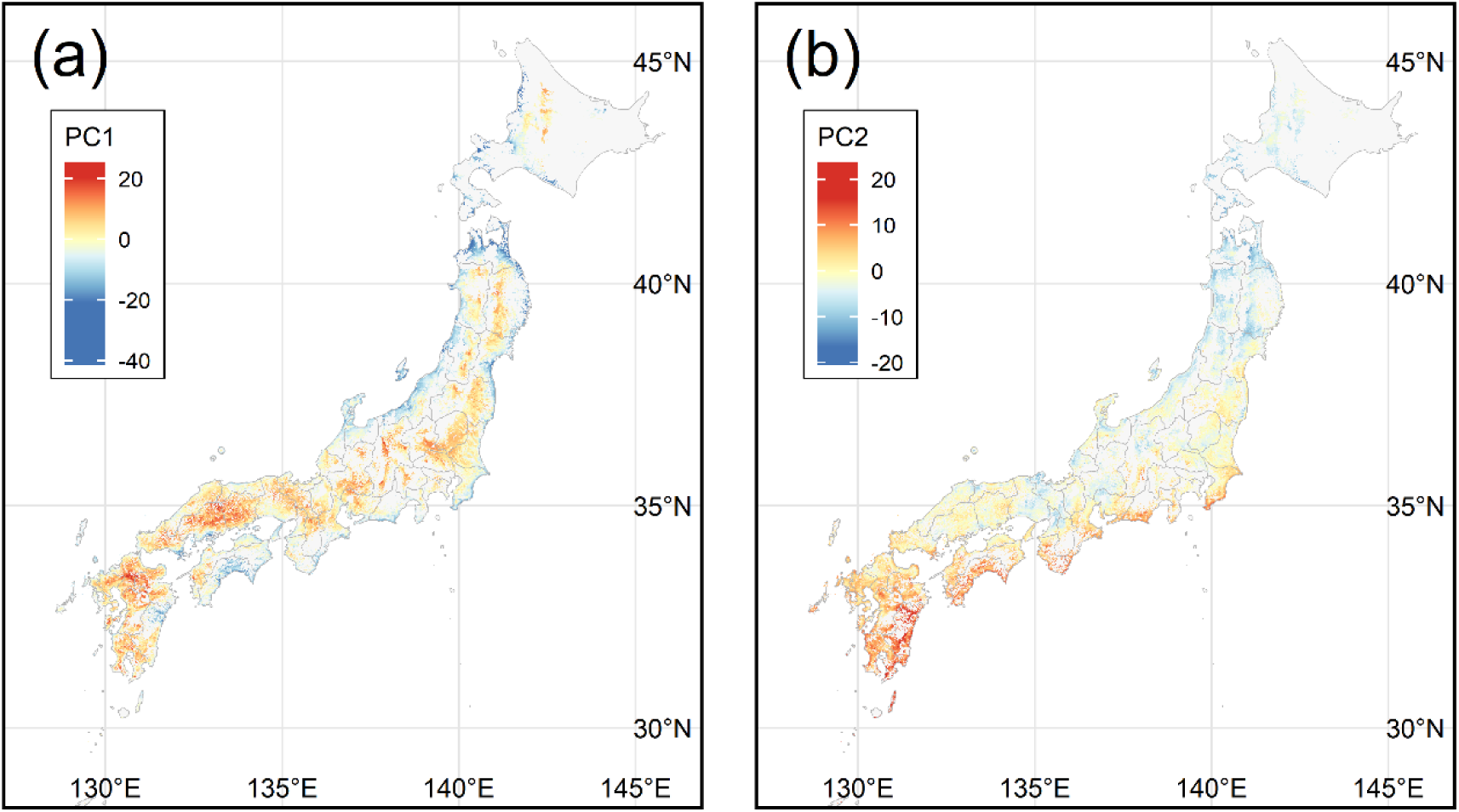
Spatial characterization of (a) PC1 and (b) PC2 scores of seasonal change in DPAT.

## Discussion

### Panicle temperatures reveal increasing HEF in Japan

Our HEF analysis, evaluated through panicle temperatures, revealed an increasing trend over time in many locations. This trend became evident in a comparison of HEF over the most recent 15 years and 15 years several decades ago (Fig. 2a, b) and was supported by statistical tests (Fig. 2c). Notably, HEFs were unprecedentedly high in the northern half of Japan in 2023, marking it as a standout year for high temperatures. This increasing trend is expected to persist for decades owing to climate change, thereby increasing the risk of spikelet sterility and heat-induced damage in rice. Measures are urgently required to prepare for unprecedented heat exposure and safeguard stable rice production in the near future.

### DPAT spatial distribution

Panicle temperatures were found to surpass air temperatures over a wide area of Japan (Fig. 1). Additionally, regional differences in the temporal means of DPAT, ranging from −0.5℃ to 3.0℃, were observed (Fig. 4b). This confirms the importance of factoring in panicle temperature, as highlighted in previous studies on spikelet sterility (Yoshimoto et al., 2011; Julia and Dingkuhn, 2013; Oort et al. 2014). Relying solely on air temperature for spikelet sterility estimation may introduce a negative bias. Furthermore, our findings reveal substantial spatial variations in DPAT seasonal changes (Figs. 6 and 7). The primary sources of variation in DPAT seasonal change were the overall magnitude of the seasonal change (PC1) and the position of peak DPAT (PC2). Based on these results, a thorough consideration of DPAT is recommended when assessing rice heat exposure, especially in areas with low altitudes where DPAT and its seasonal change tend to be more pronounced (refer to Fig. 4a for DPAT temporal mean and Fig. 8a for DPAT seasonal change). Additionally, when adjusting cropping seasons to mitigate spikelet sterility effectively, differences in the position of the peak DPAT should be considered. The positive DPAT values in Japan (ranging from −0.5℃ to 3.0℃) revealed in this study are attributed to the hot and humid weather conditions in the country. Comparable weather conditions in China have been reported to yield DPAT values reaching 4℃ (Tian et al., 2015). In contrast, in Australia, characterized by high temperatures but dry air, DPAT can fall as low as −6.8℃ (Matsui et al., 2007). Measurement and analysis of panicle temperature in 11 countries and regions revealed that regional differences in meteorological conditions led to substantial variations in DPAT among diverse climates (Yoshimoto et al., 2022). Therefore, careful consideration of meteorological differences is required when extrapolating DPAT insights from the present study to countries beyond Japan.

### Meteorological factors affecting DPAT

We analyzed the contributions of various meteorological factors to DPAT, with the results indicating that the effects of each meteorological factor could be characterized as follows: relative humidity influenced the spatial heterogeneity of DPAT, solar radiation affected the seasonal change of DPAT, wind speed exceeding 2 m s^−1^ exerted a strong cooling effect, and high maximum temperature reduced DPAT. Each characteristic is discussed in detail below.

High relative humidity was shown to increase DPAT (Fig. 4c), aligning with findings from previous studies (Matsui et al, 1997; Tian et al, 2010; Oort et al, 2014; Yoshimoto et al, 2021). Notably, relative humidity played an important role in the regional heterogeneity of DPAT: seasonal changes in relative humidity were minimal, and regions with lower mean relative humidity (Fig. S2b) tended to exhibit lower mean DPAT (Fig. 4b). Therefore, relative humidity is a pivotal factor in comparing DPAT among regions rather than predicting DPAT in a specific location. In particular, areas facing the Seto Inland Sea, where seasonal winds are obstructed by the surrounding mountains, showed consistently low relative humidity throughout the year, thereby suppressing the influence of relative humidity on DPAT.

High solar radiation was found to elevate DPAT (Fig. 4c). Although the effect of solar radiation on spatial variation in DPAT was notable, its effect was more pronounced in relation to seasonal changes (Fig. 7b). Specifically, solar radiation markedly affected the first two PCs of seasonal change. The overall magnitude of the seasonal change (PC1) was largely influenced by the difference in solar radiation between summer and autumn, whereas the position of the peak DPAT (PC2) was associated with the difference in solar radiation between June and August. This can be attributed to solar angle variations and the “Baiu” (“Tsu-yu”) rainy season in Japan from June to July. Latitude determines solar angle, affecting the overall magnitude of solar radiation, and variations in the timing and intensity of Baiu affect solar radiation in June and July. The DPAT peak in June in the northern areas of Japan (Figs. 7a and 8b), where Baiu is minimal or absent, can be explained by the change in solar angle. However, in southern areas of Japan, clouds during Baiu reduce solar radiation (from mid-June to mid-July), shifting the DPAT peak to July.

Wind speed emerged as the most influential factor explaining DPAT spatial variations: slow wind speed, especially under 2 m s^−1^ (2.5 m above the ground), led to increased DPAT (Fig. 4c). Mean wind speeds tended to be stronger on islands and along coasts (Fig. S2d), indicating that moderate winds in such locations kept DPAT low. Conversely, in other regions of Japan, low wind speed under 2 m s^−1^ should be considered when monitoring heat stress on rice panicles. Wind speed also impacted PC1 of DPAT seasonal change (Fig. 7b). Given that seasonal changes in wind speed tended to be greater at lower elevations (Fig S3), they affected the magnitude of seasonal variation in DPAT. Regarding air temperature, high maximum temperatures tended to decrease DPAT, which was evident in the DPAT temporal mean (Fig. 4c) and seasonal change in August (Fig 7b). Given that DPAT represents the deviation of panicle temperature from air temperature, panicle temperature converges toward air temperature with increasing maximum temperatures. This phenomenon can be explained from a heat balance perspective: as air temperature rises, increased evapotranspiration suppresses the temperature difference between the air and panicle. Although this aspect of DPAT tendencies has not been previously reported, similar phenomena have been observed in studies on water temperature in paddy rice fields (Fukui et al., 2017) and the temperature of an ideal wet surface (Kuwagata et al., 2022).

### Panicle temperature in high-risk regions

In this section, we explore the relationships between DPAT and meteorological factors in the seven regions identified as particularly susceptible to elevated panicle temperatures. In R1–R3, identified as regions with heightened panicle and air temperatures, mean DPAT values were low (Fig. 5a) and seasonal changes were small (Fig. 6a). However, the meteorological conditions suppressing DPAT varied by region. In R1, low solar radiation suppressed DPAT, whereas in R2 and R3, high maximum temperature and low relative humidity were inhibitory factors (Fig. 5b). Given that high maximum temperature diminishes DPAT, the increase in air temperature due to climate change may not markedly elevate DPAT in R2 and R3. Conversely, in R1, with its relatively low maximum temperature, DPAT could increase further with high solar radiation. Therefore, regions such as R1 should be vigilant of potential DPAT increases, despite their current low DPAT levels.

The remaining groups exhibiting high DPAT values, considering R4 and R5 versus R6 and R7, presented distinct characteristics. Regarding R4 and R5, high relative humidity increased mean DPAT values (Fig. 5b). These areas, facing the Sea of Japan to the north, benefit from humid winds that likely maintain high relative humidity. Additionally, delaying the cropping season tends to reduce DPAT around the heading date owing to substantial seasonal changes in DPAT (Fig. 6a). Regarding R6 and R7, high solar radiation contributed to increasing mean DPAT values (Fig. 5b). This could be attributed to these areas facing the Seto Inland Sea, which experiences limited rainfall. Moreover, the seasonal variation in DPAT tended to be low, especially in R7 (Fig. 6a). In R6, DPAT exhibited a slight decrease in September, a change which was not captured in the PCs of DPAT.

This discussion indicates that strategies for mitigating the influence of high DPAT differ by region. Given the substantial seasonal changes in DPAT in R4–R6, adjusting cropping seasons could suppress DPAT and reduce spikelet sterility. However, in R1–R3 and R7, altering cropping seasons will have little impact on DPAT. Although seasonal changes in air temperature exist, the effect of moving cropping seasons on suppressing spikelet sterility is expected to be limited compared with other regions.

## Conclusion

It has become evident that the temporal mean and seasonal change of DPAT exhibit marked differences contingent upon regional variations in meteorological conditions. Although limited to Japan, this study represents the first comprehensive analysis of DPAT trends at such a high spatial resolution. The results emphasize the need to consider multiple meteorological conditions when attempting to understand heat-induced spikelet sterility. This study shows that urgent measures are required to decrease heat exposure. For farmers, this study offers insights into the meteorological conditions leading to elevated DPAT on their farms. Agronomists and breeders can leverage the spatial and temporal heterogeneity of DPAT revealed in this study to design novel cultivation strategies and identify the requisite level of heat tolerance for future breeding efforts.

However, for a more accurate evaluation and prediction of spikelet sterility, an analysis combining actual spikelet sterility data is required. Despite previous studies on the relationship between panicle temperature and spikelet sterility, the prediction error remains substantial. Comprehending the heat stress mechanism will require rapid phenotyping techniques and a network for collecting data on spikelet sterility across a broad geographical area. The integration of model-based studies with actual phenotype data will enable a more robust and precise elucidation of this heat stress mechanism.

## Acknowledgments

This work was supported by JST-Mirai Program Grant Number JPMJMI22I2, Japan. DM was partly funded by the projects JLC-INRAE-NARO and Meta-program INRAE XRisques.

## Author contributions

YI, MY, and TK constructed a dataset of meteorological factors and panicle temperature. TT constructed a dataset of heading dates. YT conducted statistical analysis and wrote the manuscript. DM and HI supervised the analysis and edited the manuscript. All authors have reviewed and approved the final manuscript.

## Supplementary tables and figures

**Supplementary Table 1.**
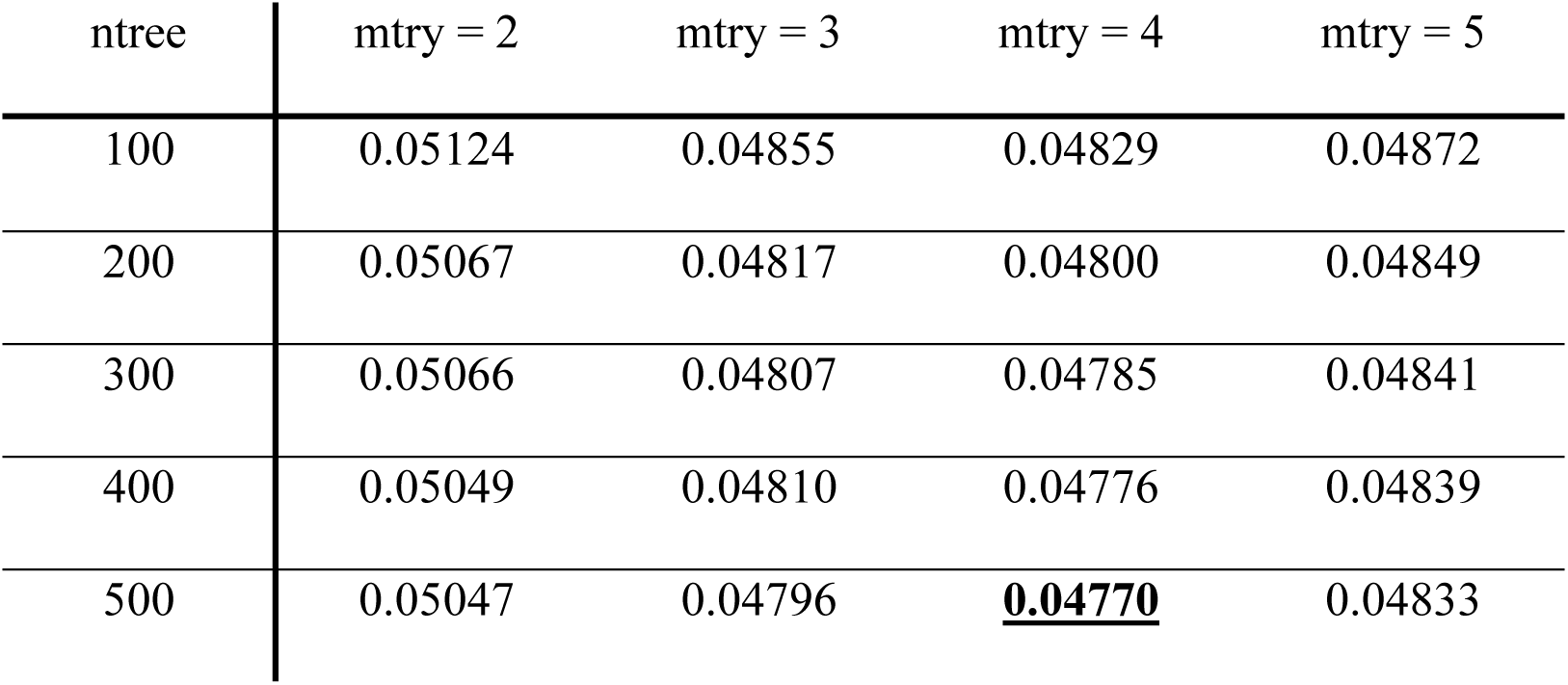
RMSE for predicting mean DPAT using a random forest model with different hyperparameters. “ntree” and “mtry” indicate the number of decision trees and the number of explanatory variables randomly selected for each tree, respectively. Six meteorological variables were used as explanatory variables. The lowest RMSE is underlined in bold.

| n tree | m try = 2 | m try = 3 | m try = 4 | m try = 5 |
| --- | --- | --- | --- | --- |
| 100 | 0.05124 | 0.04855 | 0.04829 | 0.04872 |
| 200 | 0.05067 | 0.04817 | 0.04800 | 0.04849 |
| 300 | 0.05066 | 0.04807 | 0.04785 | 0.04841 |
| 400 | 0.05049 | 0.04810 | 0.04776 | 0.04839 |
| 500 | 0.05047 | 0.04796 | **<u>0.04770</u>** | 0.04833 |

**Supplementary Table 2.** Detailed information regarding the high-risk regions. Rice field ratio: the ratio of rice field area to land area. Heat frequency: the frequency of panicle temperature at flowering time exceeding 33℃ within a year. Heat frequencies for each region were calculated as the mean heat frequencies of grid cells weighted by the rice field ratio of each cell.

| High-risk region | Land area | Rice field area | Rice field ratio | Min. heat freq. | Mean heat freq. | Max. heat freq. |
| --- | --- | --- | --- | --- | --- | --- |
| R1 | 2523 km<sup>2</sup> | 933 km<sup>2</sup> | 37% | 0.0% | 28.0% | 55.7% |
| R2 | 863 km<sup>2</sup> | 247 km<sup>2</sup> | 29% | 1.4% | 23.9% | 61.0% |
| R3 | 314 km<sup>2</sup> | 74 km<sup>2</sup> | 24% | 0.0% | 24.7% | 74.6% |
| R4 | 1077 km<sup>2</sup> | 525 km<sup>2</sup> | 49% | 0.1% | 19.0% | 50.9% |
| R5 | 1081 km<sup>2</sup> | 572 km<sup>2</sup> | 53% | 0.0% | 21.2% | 49.8% |
| R6 | 376 km<sup>2</sup> | 152 km<sup>2</sup> | 40% | 0.1% | 23.4% | 60.0% |
| R7 | 805 km<sup>2</sup> | 255 km<sup>2</sup> | 32% | 0.0% | 20.5% | 60.0% |

**Supplementary Figure 1.**
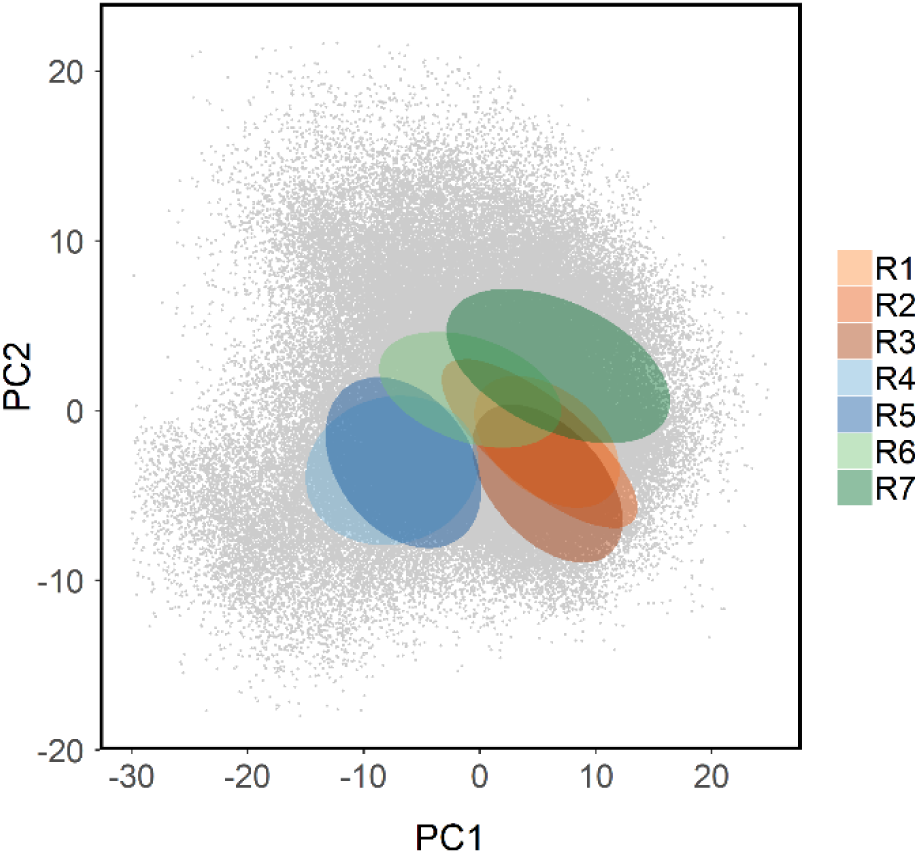
PC score distributions for the high-risk regions. Ellipses indicate the ranges in which 90% of points exist, assuming normal distributions for PC scores in each region.

**Supplementary Figure 2.**
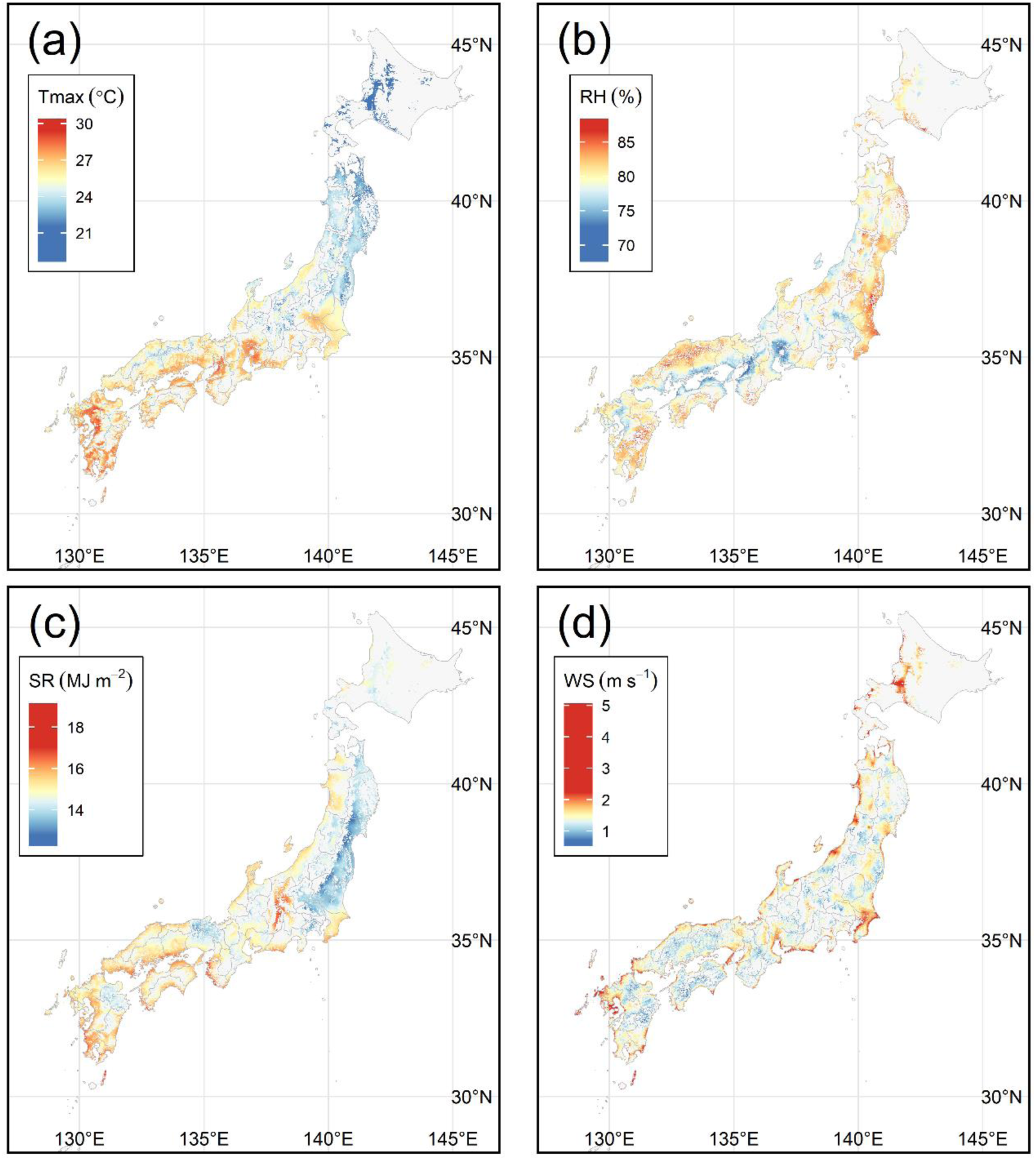
Spatial distribution of mean meteorological factors. Color bars were adjusted to ensure that their centers corresponded to the median values of the data. Results are shown for four factors: (a) maximum temperature, Tmax (℃); (b) relative humidity, RH (%); (c) solar radiation, SR (MJ m−2); and (d) wind speed, WS (m s^−1^).

**Supplementary Figure 3.**
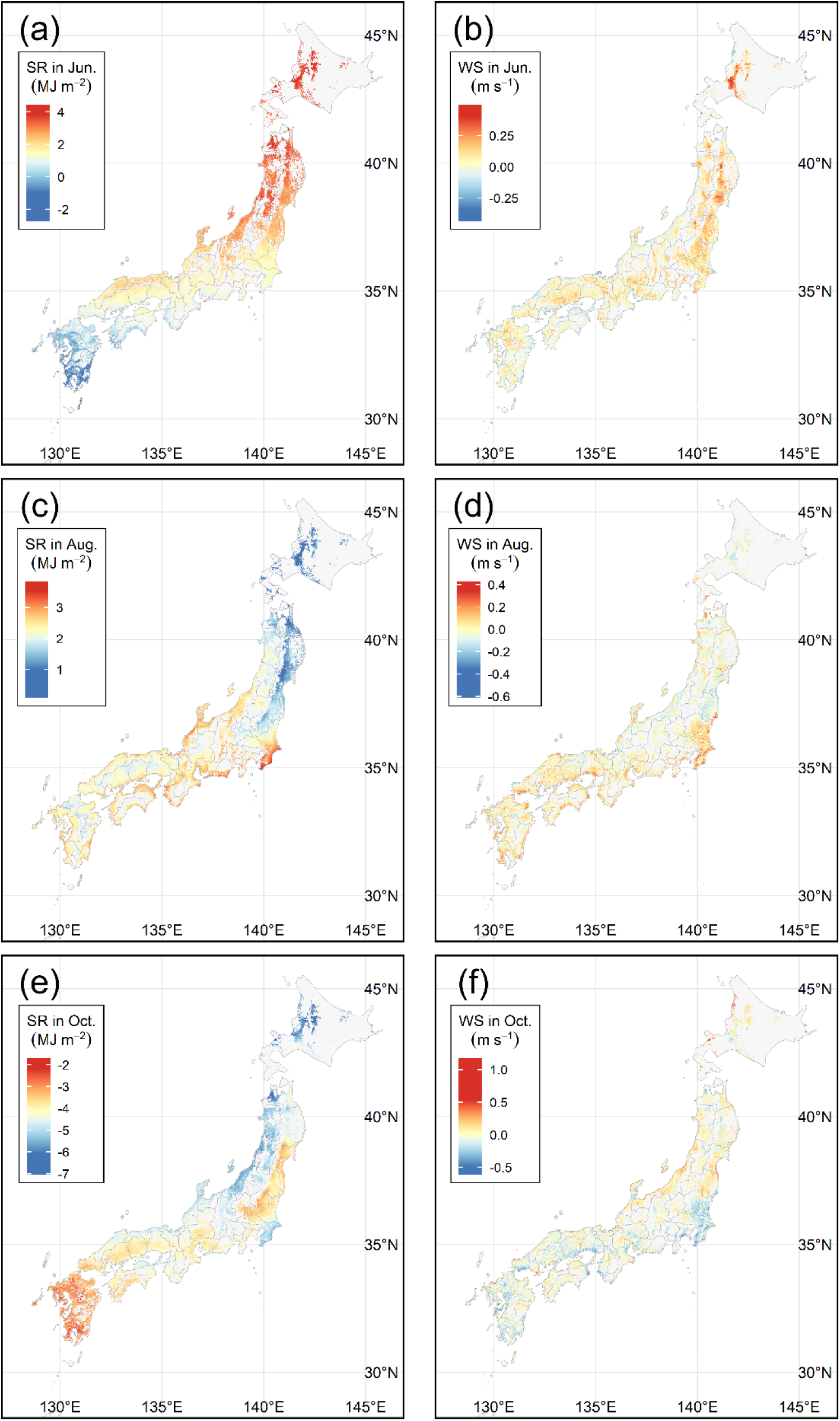
Spatial distribution of the seasonal effects of solar radiation (SR) and wind speed (WS). Color bars were adjusted to ensure that their centers corresponded to the median values of the data. Monthly means of (a, b) June, (c, d) August, and (e, f) October are shown.

